# Nonclassical nucleation of protein mesocrystals via oriented attachment

**DOI:** 10.1101/2020.08.27.267013

**Authors:** Alexander E.S. Van Driessche, Nani Van Gerven, Rick R.M. Joosten, Wai Li Ling, Maria Bacia, Guy Schoehn, Nico A.J.M. Sommerdijk, Mike Sleutel

## Abstract

Self-assembly of proteins holds great promise for the bottom-up design and production of synthetic biomaterials. In conventional approaches, designer proteins are pre-programmed with specific recognition sites that drive the association process towards a desired organized state. Although proven effective, this approach poses restrictions on the complexity and material properties of the end-state. An alternative, hierarchical approach that has found wide adoption for inorganic systems, relies on the production of crystalline nanoparticles which in turn become the building blocks of a next-level assembly process driven by oriented attachment (OA). As it stands, OA has not been observed for proteins. Here we employ cryoEM in the high nucleation rate limit of protein crystals and map the self-assembly route at molecular resolution. We observe the initial formation of facetted nanocrystals that merge lattices by means of OA alignment well before contact is made, satisfying non-trivial symmetry rules in the process. The OA mechanism yields crystal morphologies that are not attainable through conventional crystallization routes. Based on these insights we revisit a system of protein crystallization that has long been classified as non-classical, but our data is in direct conflict with that conclusion supporting a classical mechanism that implicates OA. These observations raise further questions about past conclusions for other proteins and illustrate the importance of maturation stages after primary nucleation has taken place.

## Introduction

The simplest model of crystallization assumes that the seeds formed in the initial stages are structurally identical to the macroscopic crystals that spawn from these nuclei. This idea is the foundation of the classical nucleation theory (CNT)^1^ and was until recently considered to be the most effective framework for nucleation^2^. This started to change when theoretical and experimental tools were developed that allowed scrutinizing the underlying assumptions of CNT^3^. Such enquiries to the nanoscopic mechanisms of nucleation have uncovered a broad range of nucleation pathways that do not fit the simple textbook CNT picture^4–7^ which has led to the proposal of several non-classical models of nucleation that relax the structural restraints of the nucleus^8^.

Of relevance to proteins as nucleating species is the so-called two-step nucleation model^9^ where molecules first self-organize into a metastable, liquid-like aggregate which transforms into a crystalline cluster by a structural reorganization process^9^. This model was initially formulated based on numerical simulation results, and later supported by (in)direct experimental evidence. As it stands, two-step nucleation has emerged as the consensus in the field^15,16^, but this fact remains difficult to reconcile with several issues. First, there are recorded instances of crystal nucleation where protein molecules follow a nucleation pathway akin to the one laid out by CNT^5,17,18^. Secondly, the frequently proposed candidates for the non-crystalline precursor phase of the two-step model are submicron sized particles that have been observed for numerous proteins ^14,19–23^. However, the structural nature and composition of these particles remain unknown. Their liquid-like nature has been suggested based on their propensity to flawlessly merge with the lattice of a mother crystal^12,19,24^, but that argument goes by on the lessons learned from oriented attachment of inorganic nanocrystals^25^. Thirdly, the origin of the mesoscopic size of said particles is still poorly understood. There is an ongoing debate regarding the theoretical viability of the mechanism that stabilizes their size^26–29^.

Even if one disregards these issues, the two-step model focuses narrowly on the initial stages of nucleation up until the formation of a crystalline cluster but makes no predictions regarding any later stages that may follow. To address these unknowns, we target a regime of nucleation where interactions between nuclei are likely to occur. For this we work with glucose isomerase (GI) whose nucleation mechanism has been suggested to follow a two-step pathway^19^, and for which groundwork on the characterization of the pre-nucleation particles has already been performed.

Our *in situ* data for a point mutant of GI exposes crucial interactions between nuclei mediated by OA that determine the material properties of the final phase. Moreover, we identify the prenucleation species in the I222 pathway to be nanocrystals that are hampered towards further growth due to kinetic barriers. These observations highlight the underappreciated role of the stages after primary nucleation has taken place and the impact of OA on these processes.

## Results

### GI nanocrystals merge lattices through oriented attachment

We use cryo-electron transmission microscopy (cryo-EM) to follow the nucleation of a point mutant of GI (R387A) where the surface exposed residue arginine 387 is changed to an alanine. We show that R387A crystallizes in both the I222 and H32 space group (Supporting Fig.1). H32 exhibits high nucleation rates which increases the probability for nanocrystal interactions to occur in solution. Almost immediately after mixing R387A with PEG 1000, we can resolve nano-crystalline particles (Fig.1a, 1min40). These nanocrystals are facetted and their FFTs show clear diffraction patterns that match the predictions based on crystallographic data obtained from macroscopic crystals (a=b=6.64nm, 60°; Supporting Fig.2). On some occasions, we measure minor stretching of these intermolecular distances (6.7-6.9nm) indicating that there is some flexibility in these nanocrystal lattices. We also point out that the facets, although rough on some occasions, are surprisingly smooth, suggesting that they represent a Wulff shape that emerges out of the anisotropy in the surface tension. Moreover, the hexagonal shape of the nanocrystals is in line with what can be expected from a simple periodic bond chain analysis for the H32 space group.

**Fig.1:**
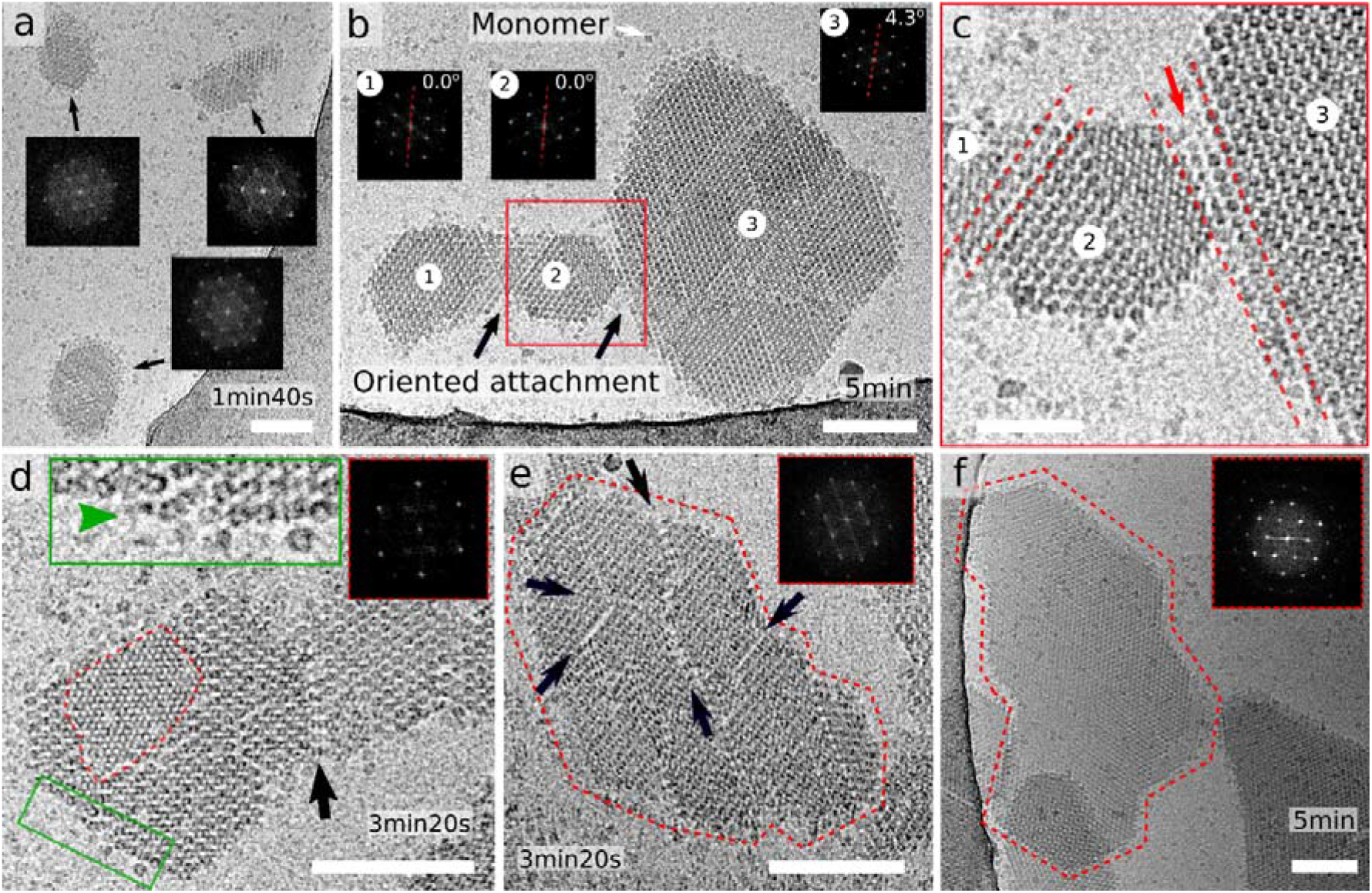
Oriented attachment of GI R387A nanocrystals: (a) submicron nanocrystals formed 1min40s after mixing protein and precipitant; corresponding FFT images exhibit sharp maxima; (b) oriented attachment of individually nucleated nanocrystals into a larger, merged lattice composed of domains 1 and 2, (c) making loose lateral contact with domain 3 at an angle of 4.3°; (d) two nanocrystals with merged lattices; red inset: FFT of the second molecular layer that is forming on the parent crystal; green inset: zoom-in of the unfinished molecular layer resolving local disorder and incoming growth units; (e) and (f) Large aligned nanocrystal assemblies with and without fault lines between the separate domain, respectively. Panels (b) - (f) were recorded from aliquots plunge frozen after 3-5min incubation. Scalebar is 100nm in panels a,b,d,e and f, and 50nm in c.

**Fig.2:**
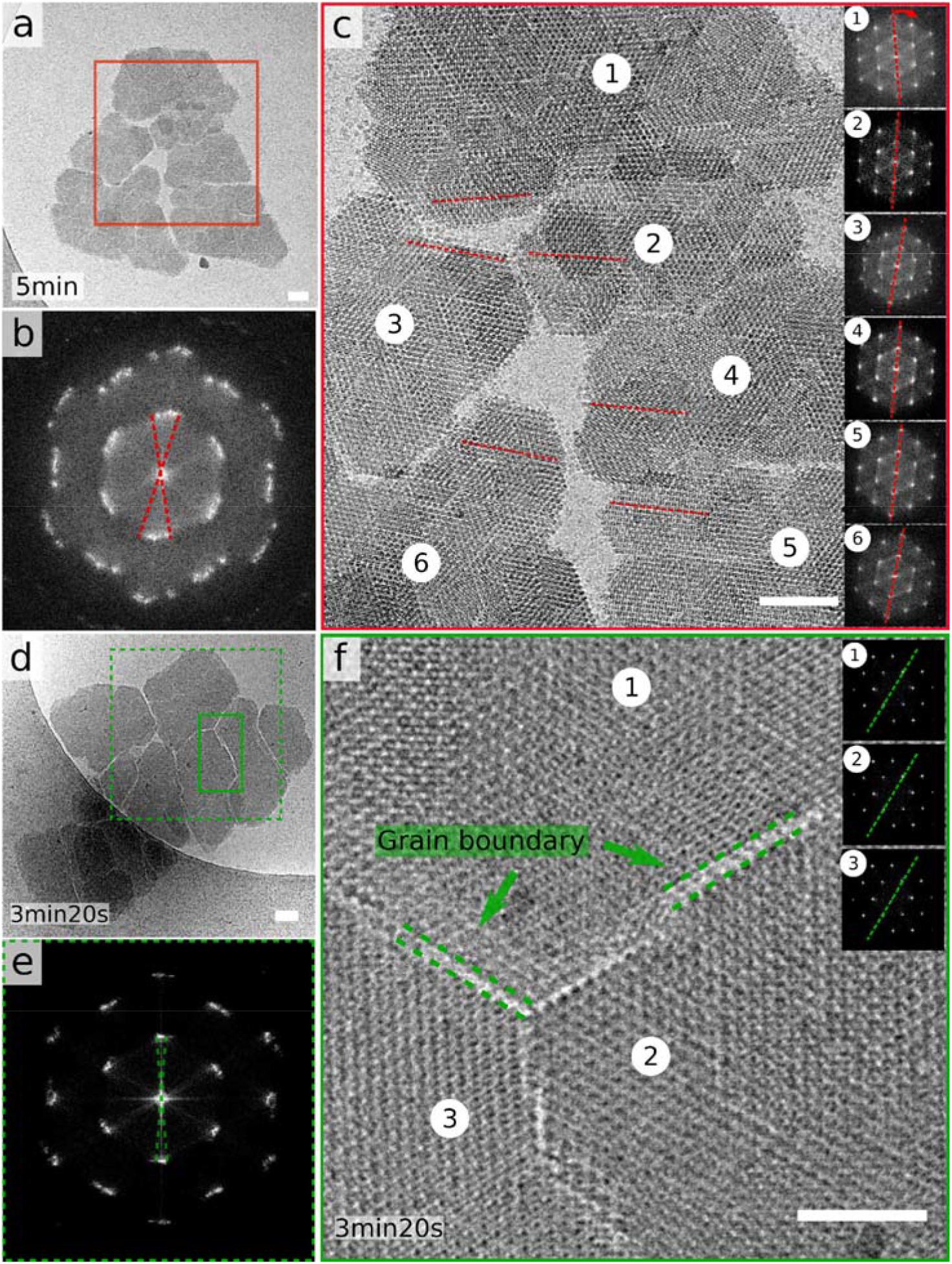
GI R387A mesocrystals: (a), (c) and (d) large micron-sized composite nanocrystal structures with pronounced fault lines that separate lattice domains that are in near-alignment as demonstrated by their respective FFTs (b) and (e); zoom-in of the grain boundaries that span one or two molecular distances between the individual domains (f). The inset in (f) demonstrates the high degree of lattice order within each separate domain: domain 2 shown as representative example. Scalebar is 50nm in panels a,c and d, and 25nm in f.

At later time points of the crystallization reaction (3min), we find groupings of similar nanocrystals that have merged into a unified lattice with no discernable stacking faults at their junctions (Fig.1b-d, 5min). The crystal in Fig.1b is of interest because it shows two smaller domains (1 and 2) that are in perfect co-alignment, and that make loose contact with a larger, third domain sitting at a 4.3° angle. It is our assertion that such conglomerate structures are formed through an oriented attachment process. In fact, close inspection of the junction area between domain 2 and 3 reveals that both regions are in near contact with each other (Fig.1c). It is tempting to speculate that the angular offset between these domains would eventually be eliminated by rotational diffusion, leading to the melding of both lattices once alignment had been reached^33^. A similar event is shown in Fig.1d (and Supporting Figure 3), where we discern two nanocrystals that have melded in coalignment into one unified lattice, held together by a joint molecular column at their interface (black arrow Fig.1d and white arrow Supporting Figure 3c).

**Fig.3:**
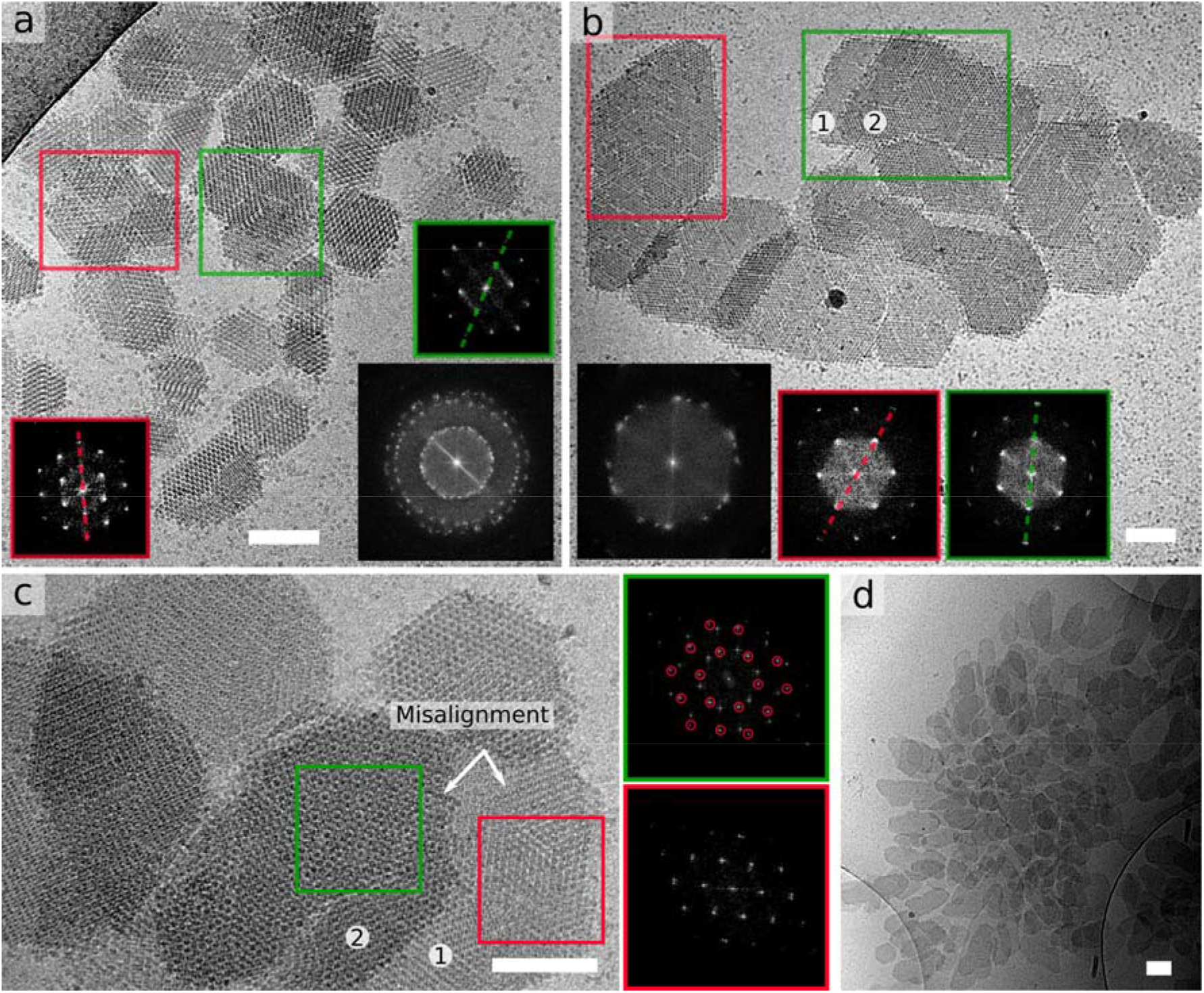
Lack of inter-crystal alignment in large nanocrystal assemblies: (a) poly-crystalline cluster with local hotspots of alignment (see FFT insets), (b) example of two vertically stacked domains exhibiting registry between their corresponding lattices (blue) but a 26° misalignment with the region enclosed in pink, (c) example of lack of axial alignment between two nanocrystals where the FFT of the interlaced pattern (green) reveals two independent lattices residing at an angle of 25°, (d) disordered grouping of over 100 nanocrystals. Scalebar is 100nm.

A mechanism of cooperative, and guided merging of separately nucleated lattices is implied by observations of large (~0.5μm) more erratically shaped structures that deviate from the expected Wulff shape and which display faint internal contrast lines that outline the individual clusters residing within the larger lattice (black arrows Fig.1e). These lines are either grain boundaries (GB) and therefore represent local points of failure to completely merge lattices, or they are thin layers of solvent that still need to be expelled for contact to be completed. Successful docking of multiple nanocrystals is certainly possible as evidenced by the complex morphology in Fig.1f that does not appear to have any discernable GBs.

We find even larger (>1μm) composite structures with pronounced fault lines that separate homogeneous lattice domains (Fig.2a and d). What is striking is that all these domains are in near perfect alignment, which means that these superstructures are (near) mesocrystals. This can be inferred from the FFT (Fig.3b) of the highlighted region in Fig.2a showing only minimal spread of the diffraction peaks. In panel c of Fig.2 we also show the individual FFTs of the regions 1 to 6, highlighting the various degrees of rotation. The mesocrystal in Fig.2d exhibits even better alignment of its subdomains as demonstrated by Fig.2e. Careful analysis, though, of the orientations of domains 1, 2 and 3 taking domain 1 as reference, learns that they reside at angles 0 ± 0.4°, -0.8 ± 0.1° and 2 ± 0.3° of each other. Moreover, the typical distance across the GBs is in the range of one or two molecular rows, suggesting that the shape complementarity of the various domains is the result of the addition of new GI molecules at the imperfect contact areas that form after docking (Fig.2f).

We do, however, also find examples where the inter-crystal alignment has either failed or has not yet reached completion before cryo-quenching took place, resulting in an overall lack of the long-range order (Fig.3). We identify an absence of lattice alignment between nanocrystals that interact laterally or stack axially (Fig.3a-c). For example, the green area in Fig.3b highlights two stacked domains exhibiting registry between their corresponding lattices, but clear misalignment with the region enclosed in red. Fig.3c is an example of lack of axial alignment where the FFT of the interlaced pattern (green) reveals two independent lattices residing at an angle of 25±0.5°. Although cryo-EM images represent single snapshots in time, it is tantalizing to speculate that these structures may eventually relax into a lower energy state possessing more long-range order. There may, however, be a limit to this self-organization process. It seems unlikely that the disordered grouping of over 100 nanocrystals in Fig.3d will transform into a single, unified (meso)crystal. Indeed, light microscopy observations of the earliest observable crystals appear twinned and full of growth defects (Supporting Fig.4) reminiscent of the alignment failures we see at the nanoscopic level.

**Fig.4:**
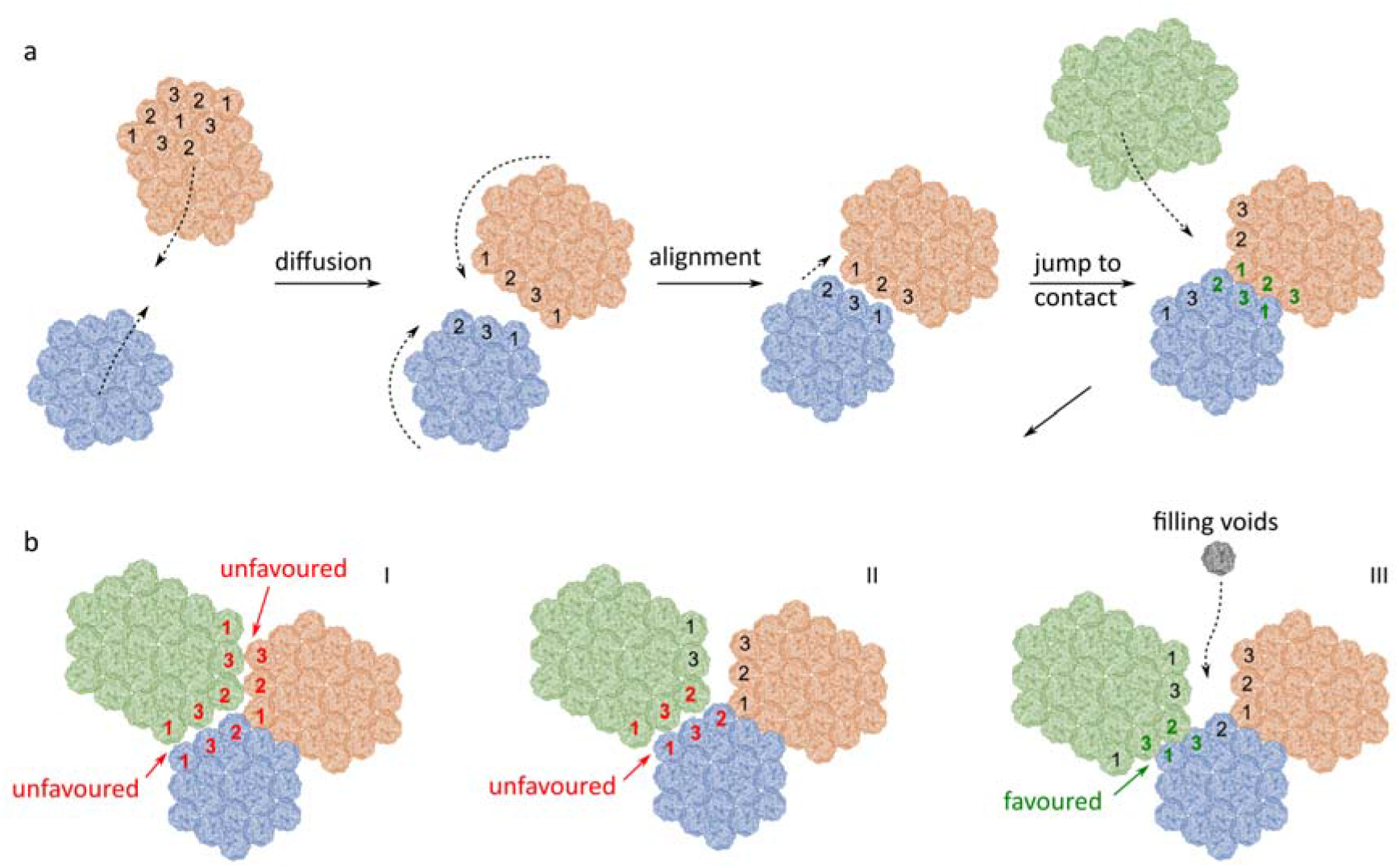
Model for OA of GI nanocrystals: GI molecules pack along a 3-fold axis within the (001) plane from which we discern three different GI orientations (1,2,3). Nearest neighbors (NN) exclude GI molecules with identical orientations; (a) Simplified scheme of self-assembly: freely diffusing nanocrystals approach each other, followed by rotational and translational adjustments to align both lattices. Alignment facilitates a final jump to contact by desolvation of the surface patches that partake in lattice contact formation; (b) Illustration of three different scenarios for further growth: I and II violate H32 symmetry rules and are likely to lead to the formation of a GB at the interface; III leads to successful merger of all 3 lattices and the resulting voids can be filled by monomer addition.

### Rotational symmetry restraints limit the probability of success for the final jump to contact

The OA mediated accretion of R387A nanocrystals is remarkable considering the non-trivial symmetry requirements of the H32 space group (Supporting table 1 and Supporting Discussion). Based on the crystal structure of the planar H32 crystals observed in cryo-EM, we identify the (001) plane as the dominant orientation. The 3-fold screw axis is perpendicular to the (001) plane resulting in 3 distinct GI orientations (designated as 1, 2 and 3 in Fig.4a). The nature of the crystallographic symmetry is such that identically oriented GI molecules do not engage in lattice contacts. This poses additional registry requirements for OA to be successful, as highlighted in Fig.4b. Even if two proximate nanocrystals are in rotational alignment, approximately only a third of all configurations will lead to proper docking of both lattices (Supporting Figure 5). For those scenarios where bond formation is not possible, crystals will either need to slide laterally along the interface or diffuse away to reinitialize altogether. We use PEG 1000 as a crystallization agent that induces an attractive depletion force that scales with the size of the interface between crystallites as they approach each other. Hence, for larger clusters it may become increasingly unlikely to find the proper translational register as a result of the depletion force even tough rotational alignment has already been reached. This fits with the cryo-EM observations, which show that grain boundaries tend to develop within larger conglomerates.

**Fig.5:**
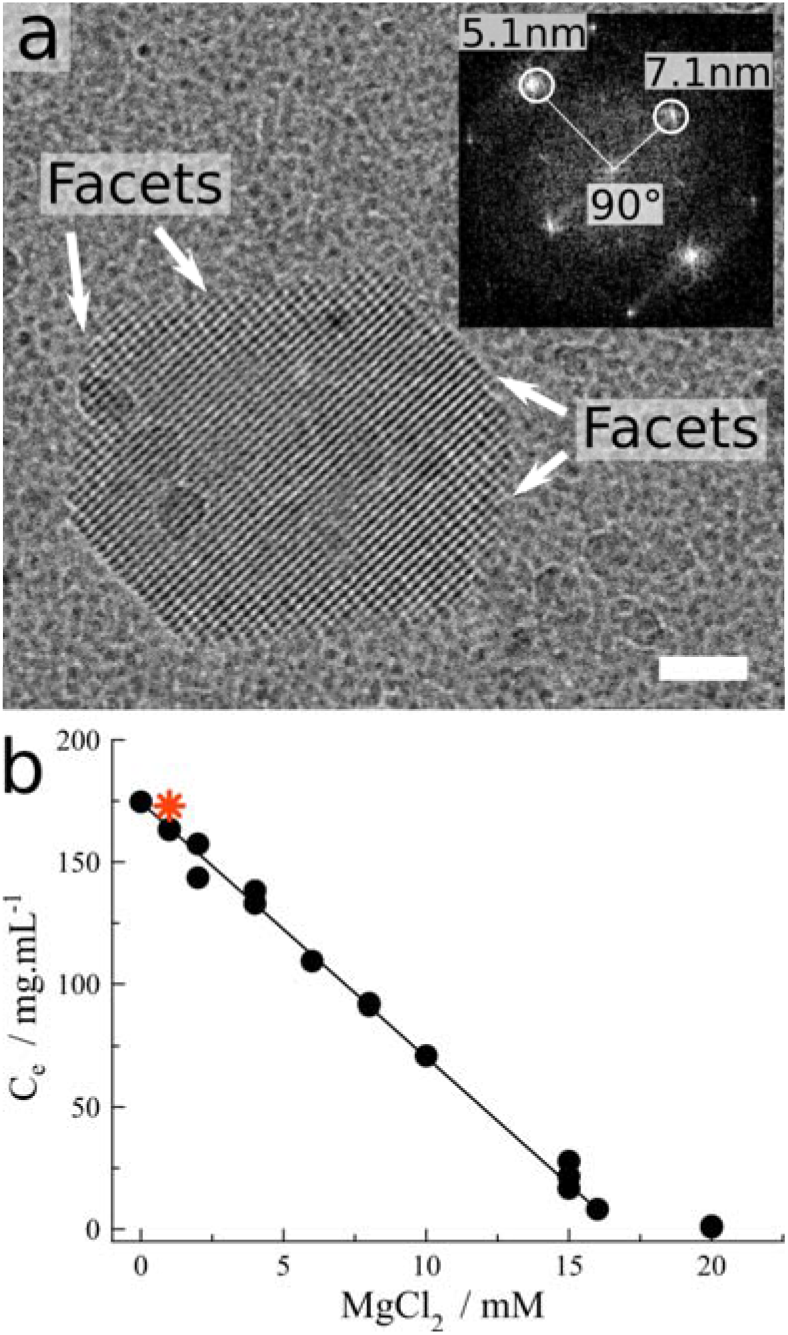
(a) cryoTEM image a facetted nanocrystal in the unfiltered GI stock solution, with lattice fringes visible along two directions; inset: FFT of the central region of the crystal exhibiting clear maxima that correspond with the experimentally measured distances between the lattice planes of the I222 space group. Scalebar is 50nm; (b) Solubility curve of orthorhombic GI crystals as a function of MgCl_2_ concentration. Line is a guide for the eye.

### OA may be a universal mechanism for proteins

To the best of our knowledge there are no direct observations of OA for bio-macromolecular systems, and R387A represents the first example for this class of molecules. At the same time, there is a host of experimental observations that remain enigmatic but that can be resolved if OA is implicated. We refer to mesoscopic protein particles merging with the lattice of a mother crystal in a largely defect-free manner^12,19,24^. The remarkable efficiency by which this happens has prompted us and others in the past to postulate that said particles are liquid-like in nature, and that these particles undergo a solidification transformation upon contact, accommodating the lattice of the mother crystal in the process. That interpretation was favored because the probability for a protein nanocrystal to rotate towards perfect registry with another lattice through random Brownian motion was considered negligible^12,19,30^. Using cryo-EM we are now in a position to structurally characterize these elusive particles that have been put forth as the intermediary species in the two-step nucleation pathway. Extensive cryo-EM screening of plunge-frozen aliquots of an unfiltered, wild type GI stock solution reveal the presence of imperfect rhombic nanoparticles with clear lattice fringes. They measure 300 ± 150 nm across (n=12; Fig.5b, Supporting Fig.6) which fits the published size range of the pre-nucleation particles for GI^19^. The intermolecular distances (5.1nm and 7.1nm at 90°) derived from the maxima in the calculated fast Fourier transform (Fig.1c) are in line with previous observations for I222 GI crystals and fit predictions based on X-ray crystallographic data^5^. This readily explains the reported seeding behavior at the macroscopic level, as well as the specificity regarding the I222 polymorph. Based on solubility measurements we conclude that our concentrated protein stock solution is slightly supersaturated (6%) with respect to crystalline phase. Using previously determined kinetic parameters we can estimate their expected crystal growth rate (see Supporting Discussion). We predict it to be in the range of one molecular layer per day, which explains the apparent size stability of these nanocrystals in the stock solution. Moreover, we link their formation to temporary concentration gradients that form when we concentrate the freshly dialyzed protein solution prior to long-term storage (Supporting Fig. 10).

From this we conclude that particle merger events previously published for macroscopic GI crystals are likely driven by OA. This prompts a reinvestigation of other systems and suggests that OA may be a widespread mechanism for proteins.

## Discussion

Macromolecular crystallization is an astounding feat of nature. Even though proteins are large, dynamic and often highly anisotropic molecules, they can form a minimal assembly that guides incoming molecules to registry as instructed by its internal rules of symmetry. Understanding how this nucleus forms has been the subject of debate for over two decades now. The two-step nucleation model is arguably the most popular and is often considered as a consensus view on this subject. Direct experimental evidence for a nucleation trajectory akin to the predictions of the two-step model have recently been provided by Houben *et al* for the case of ferritin crystallization^11^. But the process that they witness is more nuanced than the simplified original two-step scheme. They record the initial formation of disordered ferritin aggregates that tend to increase in both order and density from their surface towards their interior in a cooperative process of gradual desolvation. The process of self-assembly that they describe is in stark contrast with our observations here for GI and it’s point mutants. Although there have been indications that GI may also first condense into a non-crystalline precursor under certain conditions^19^, our cryo-EM observations identify these particles to be nanoscopic renditions of one of the polymorphs GI can crystallize in. This essentially means that GI I222 crystals nucleate in qualitative accordance with classical predictions. Based on our previous work on GI^5,17^ and the work of others^11^, it is clear that proteins can nucleate through a number of routes that are conceptually quite diverse and which do not fit our idealized views.

The differentiation between one or two-step nucleation becomes less relevant for situations where the number of nucleation centers is relatively high, such that interactions between nuclei cannot be disregarded. R387A showcases such a regime of nucleation in the high concentration limit of nucleating entities. Here we see clear interplay between independently formed clusters in a manner reminiscent to a host of inorganic systems whereby nuclei diffuse in solution, collide, and coalesce to form larger unified lattices and mesocrystals^34^. This merger of lattices in a defect-free manner is unlikely to occur simply by chance, rather, nanoparticles experience a torque guiding each other towards (near) perfect registry^33^. That directed maneuvering has long been recognized as a widespread mechanism of nanoparticle assembly and is referred to as oriented attachment (OA)^35^. The steering torque that guides OA can often be attributed to the dipole moment of the nanocrystals^36^. This is not the case for GI because the net dipole moment of a GI tetramer, i.e. the crystals’ building block, is zero because the dipole moments of the monomers cancel each other. And yet, the GI mesocrystals strongly indicate that such a guiding mechanism exists because we see alignment of crystalline domains that do not have any bridging contact points between their respective lattices. This demonstrates that alignment occurs before docking, i.e. OA is facilitated by lattice alignment, and presumably at relatively long range (~10nm). OA for dipole-free systems has been attributed to short range Van der Waals attractions, but Liu *et al* have recently suggested that local water structuring could play a role in the long-range steering that takes place before the jump to contact^37^.

At the same time, R387A also demonstrates the limits of OA. As these crystals grow larger, GB become more pronounced and mosaicity increases. Understanding the underlying forces that contribute to the successful OA of protein nanocrystals is relevant for the bottom-up design of novel bio-materials where precise control over the self-assembly mechanism could help tune the size, aspect ratio and polycrystallinity of the final phase^38^. And with the cryoEM revolution in structural biology that is focusing more on electron diffraction^39^, combined with the need for protein nanocrystals for XFEL diffraction^40^, we believe a better understanding of macromolecular OA could contribute in these domains.

## Materials and Methods

### Protein production and purification

Glucose Isomerase (GI) was obtained from Hampton Research (wild type *Streptomyces rubiginosus*) and received as a crystalline slurry. Small aliquots were dialyzed overnight (Spectra/Por Standard RC Turbing: 12-14kDa; Spectrumlabs) against 10mM Hepes pH 7.0, 1mM MgCl_2_ at 4°C. The protein solution was concentrated using a centrifugal filter with a MWCO 100kDa (Amicon Ultra −15 Cellulose, Milipore) to 173mg·mL^1^ and stored at 4°C. Concentrations were determined by measuring the absorbance at 280nm using an extinction coefficient ε_280_ of 1.074 mg^1^mL.cm^−1^. Stock solution was used as is for cryoTEM imaging.

GI R387A was produced recombinantly as previously described^5^. In brief, R387A was expressed in *E. coli* BL21(DE3) after induction at OD_600nm_ of 0.7 with 1mM IPTG for 3 h at 37°C. Cells were harvested by centrifugation at 6238 g for 15 minutes and resuspended in 100mM Tris-HCl pH 7.3, 1mM ethylenediaminetetraacetic acid (4mL.g^−1^ wet cells) supplemented with 5μM leupeptin, 1mM 4-(2-aminoethyl)benzenesulfonyl fluoride (AEBSF), 100 μg.mL^−1^ lysozyme and 20 μg.mL^−1^ DNase I and incubated for 30min at 4°C. Subsequently, MgCl_2_ was added to a final concentration of 10mM and cells were lysed by two passages in a Constant System Cell Cracker at 20 kpsi at 4 °C and cell debris was removed by centrifugation at 48,400 g for 45 min at 4°C. The cytoplasmic extract was incubated for 10min at 65°C and the insoluble fraction was removed by centrifugation at 48,400g for 45min at 4°C. The supernatant was filtrated through a 0.22μm pore filter and loaded on a 5 ml pre-packed Hitrap Q FF column (GE Healthcare) equilibrated with buffer A (50mM bis-tris-HCl pH 6.0, 10mM NaCl). The column was then washed with 40 bed volumes of 20% buffer B (50mM bis-tris-HCl pH 6.0, 1M NaCl) and bound proteins were eluted with a linear gradient of 20–50% buffer B over 10 bed volumes. Fractions containing R387A, as determined by SDS–PAGE, were pooled and supplemented with ammonium sulfate to a final concentration of 1.5M and loaded on a 5ml pre-packed HiTrap Phenyl HP column (GE Healthcare) equilibrated with buffer A (100mM Tris pH 7.3, 1.5M ammonium sulfate). The column was then washed with 40 bed volumes of 25% buffer B (100mM Tris pH 7.3) and bound proteins were eluted with a linear gradient of 25–85% buffer B over 15 bed volumes. Fractions containing R387A were pooled and dialyzed (Spectra/Por Standard RC Turbing: 12-14kDa; Spectrumlabs) against 10mM Hepes 7.0, 1mM MgCl_2_ over night at 4°C (buffer was replaced twice) and concentrated in a 100 kDa molecular weight cutoff spin concentrator (Amicon Ultra −15 Cellulose, Milipore)to a typical final concentration of 30mg.mL^−1^.

### Glucose Isomerase crystallization

To trigger crystallization of wild type GI and R387A, the protein stock solutions were mixed at 22°C with an equal volume of 100mM Hepes 7.0, 200mM MgCl_2_ and 8% (w/v) PEG_1000_.

### Dynamic Light Scattering

Intensity correlation functions of filtered and non-filtered GI stock solutions were collected at 20°C in 4μl disposable cuvettes using a DynaPro Nanostar (Wyatt).

### Solubility measurements

Pre-grown I222 GI crystals were centrifuged (5min, 14k rpm) and washed three consecutive times using 1ml fractions of 10mM Hepes 7.0, and 0 – 20 mM MgCl_2_ before being stored at 4°C. Protein concentration determination of the soluble phase after centrifugation was done on a weekly basis until a steady-state was reached.

### Cryo-Transmission Electron Microscopy

For cryo-TEM, 200 mesh Cu grids with Quantifoil R 2/2 holey carbon films (Quantifoil Micro Tools GmbH) were used. Sample preparation was performed using an automated vitrification robot (FEI Vitrobot Mark III) for plunging in liquid ethane^41^. All TEM grids were surface plasma treated for 40 seconds using a Cressington 208 carbon coater prior to use. The samples were imaged with the TU/e cryoTITAN (FEI, www.cryotem.nl) operated at 300 kV, equipped with a field emission gun (FEG), a post-column Gatan Energy Filter (GIF) and a post-GIF 2k x 2k Gatan CCD camera. We choose *t_0_* as the moment where we induce supersaturation with respect to the crystalline phase (i.e. mixing of the protein with the precipitant solution) and *t_end_* as the time at which crystals become detectable using light microscopy. The exact time point of the samples as indicated in the main text is defined as the moment (after blotting excess liquid) when the EM grid is plunged into the liquid ethane. Images were acquired in low-dose mode at a magnification of either 24000× with a nominal defocus of −5 μm or 11500× and −10 μm defocus.

### Crystallographic Analysis

Nearest crystallographic neighbors of the GI molecule are generated using Chimera 1.13.1. Residues partaking in lattice contacts are identified by calculating the accessible surface area (ASA) on a per-residue level using AREAIMOL of the CCP4 software suite^42^. ASAs are determined for both the starting models as well as the models consisting of the GI molecule and its nearest neighbor using a probe radius of 1.4Å. Residues with a non-zero difference in accessible surface area (ΔASA) are (partially) buried in the bound complex and therefore considered to be part of the lattice contact patch. Hydrogen bond pairs are identified using the *FindHBond* tool in Chimera 1.13.1rc using default settings, and salt-bridges are identified using the PDBePISA (http://www.ebi.ac.uk/pdbe/pisa/) and the 2P2I (http://2p2idb.cnrs-mrs.fr/2p2i_inspector.html) protein interaction webservers.

## Author contributions

MS and AESVD designed the project and carried out the crystallization and light scattering experiments. NVG cloned, recombinantly expressed and purified R387A. Cryogenic freezing and cryoEM imaging was performed by WLL, MB and RRMJ. MS supervised the study. MS, AESVD and NAJM wrote the paper, with contributions from all authors.

## Acknowledgments

MS acknowledges financial support by the FWO under project G0H5316N and 1516215N. This work made use of the Center of Multiscale Electron Microscopy at the Eindhoven University of Technology (www.cryotem.nl) and the EM facility at the Grenoble Instruct-ERIC Center (ISBG; UMS 3518 CNRS CEA-UGA-EMBL) with support from the French Infrastructure for Integrated Structural Biology (FRISBI; ANR-10-INSB-05-02) and GRAL, a project of the University Grenoble Alpes graduate school (Ecoles Universitaires de Recherche) CBH-EUR-GS (ANR-17-EURE-0003) within the Grenoble Partnership for Structural Biology. The IBS Electron Microscope facility is supported by the Auvergne Rhône-Alpes Region, the Fonds Feder, the Fondation pour la Recherche Médicale, and GIS-IBiSA.

