## Supporting Information for "Nonclassical nucleation of protein mesocrystals via oriented attachment"

Contents:

Supporting Figures: 10

Supporting Table: 2

Supporting Discussion

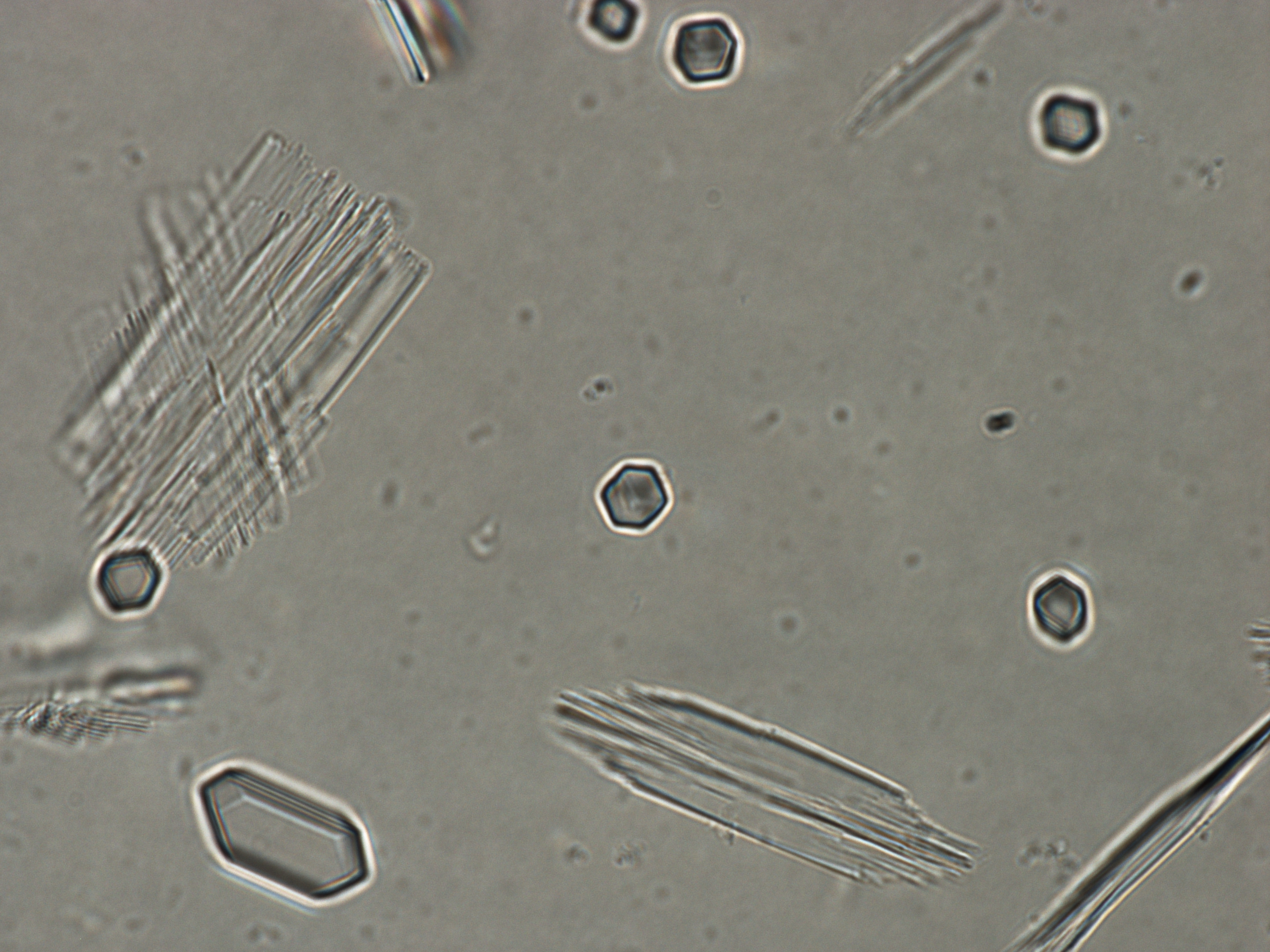

Supporting Figure 1: Polarized light microscope image of two GI R387A polymorphs (rhombic: I222; twinned platelets: H32) formed 10 minutes after mixing protein stock solution with PEG_1000_ (10min, 100x objective).

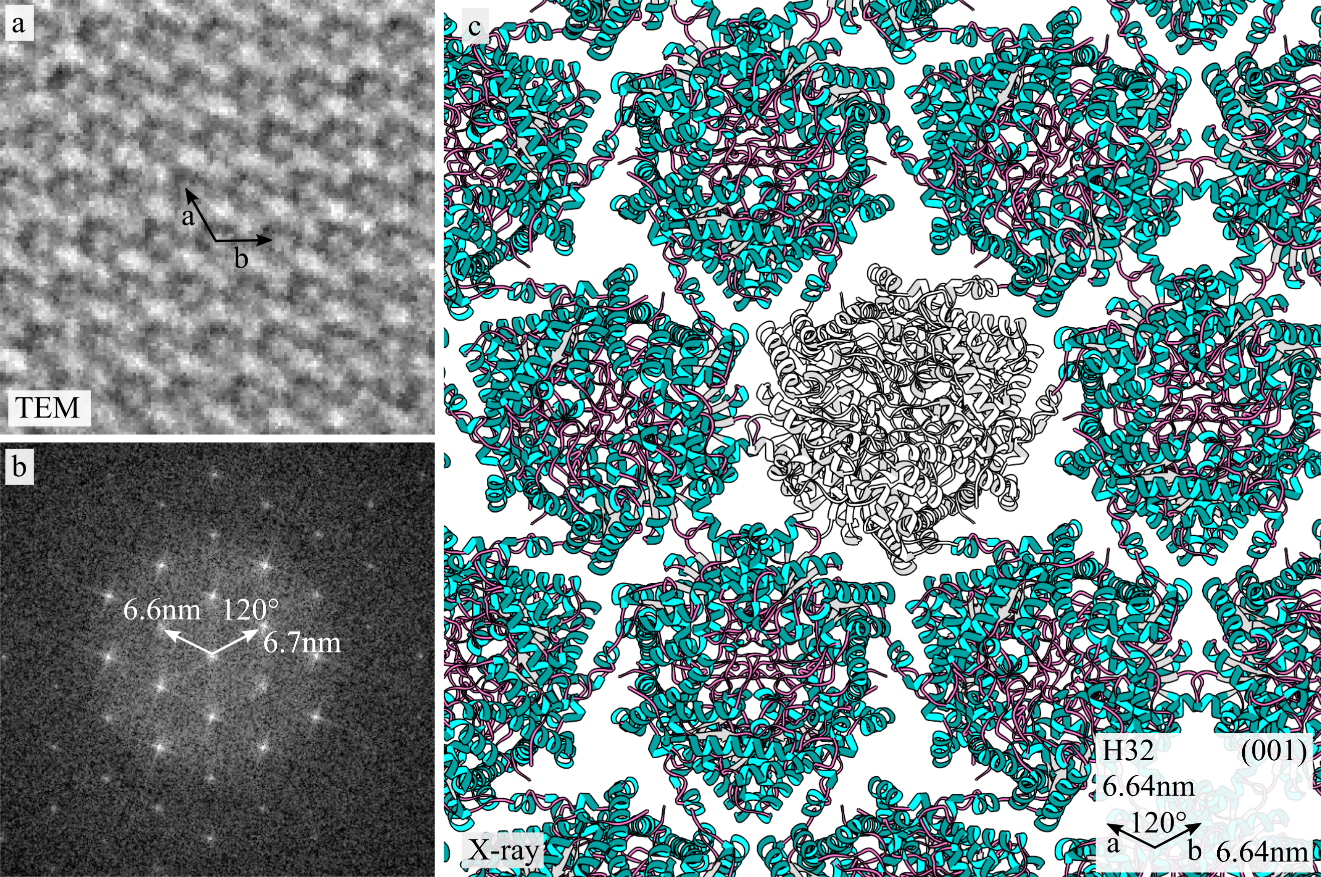

Supporting Figure 2: (a) Raw cryoTEM image of the lattice of R387A glucose isomerase nanocrystals formed in 15 mg.mL^-1^ glucose isomerase, 50mM Hepes pH 7.0, 100mM MgCl_2_, 4% (w/v) PEG 1000; (b) FFT of panel (a) with indicated nearest neighbor distances and measured angle; (c) crystallographic arrangement of glucose isomerase molecules along the (001) plane of the trigonal H32 space group (as determined by X-ray diffraction of S171W crystals) with the theoretical nearest neighbor distances along the a and b axis and corresponding angle (inset).

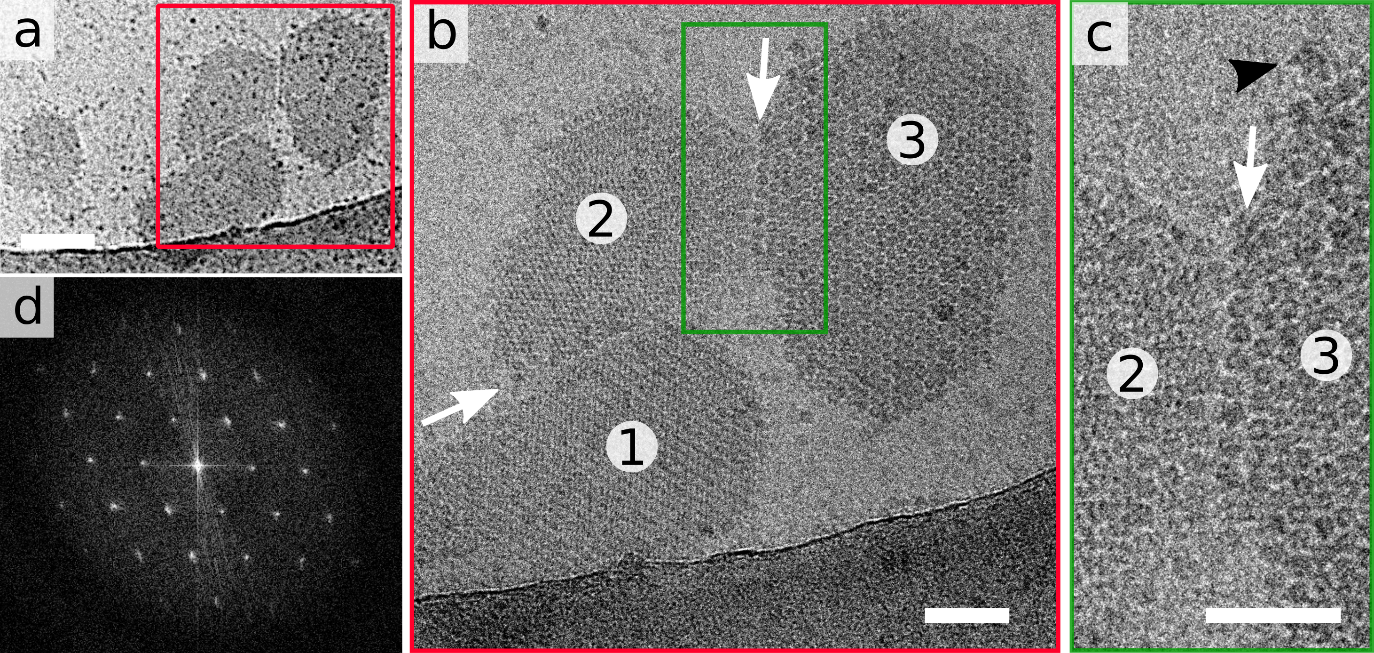

Supporting Figure 3: Additional example of OA of 3 R387A GI nanocrystals: low magnification image (a) with consecutive zoom-ins of the local cluster (b) and the binding interface between 2 and 3 encompassing approximately 8 GI molecules. Black arrowhead: incoming GI tetramer (c); the sharp maxima in the FFT of the highlighted area (red) demonstrates excellent alignment of the respective lattices (d).

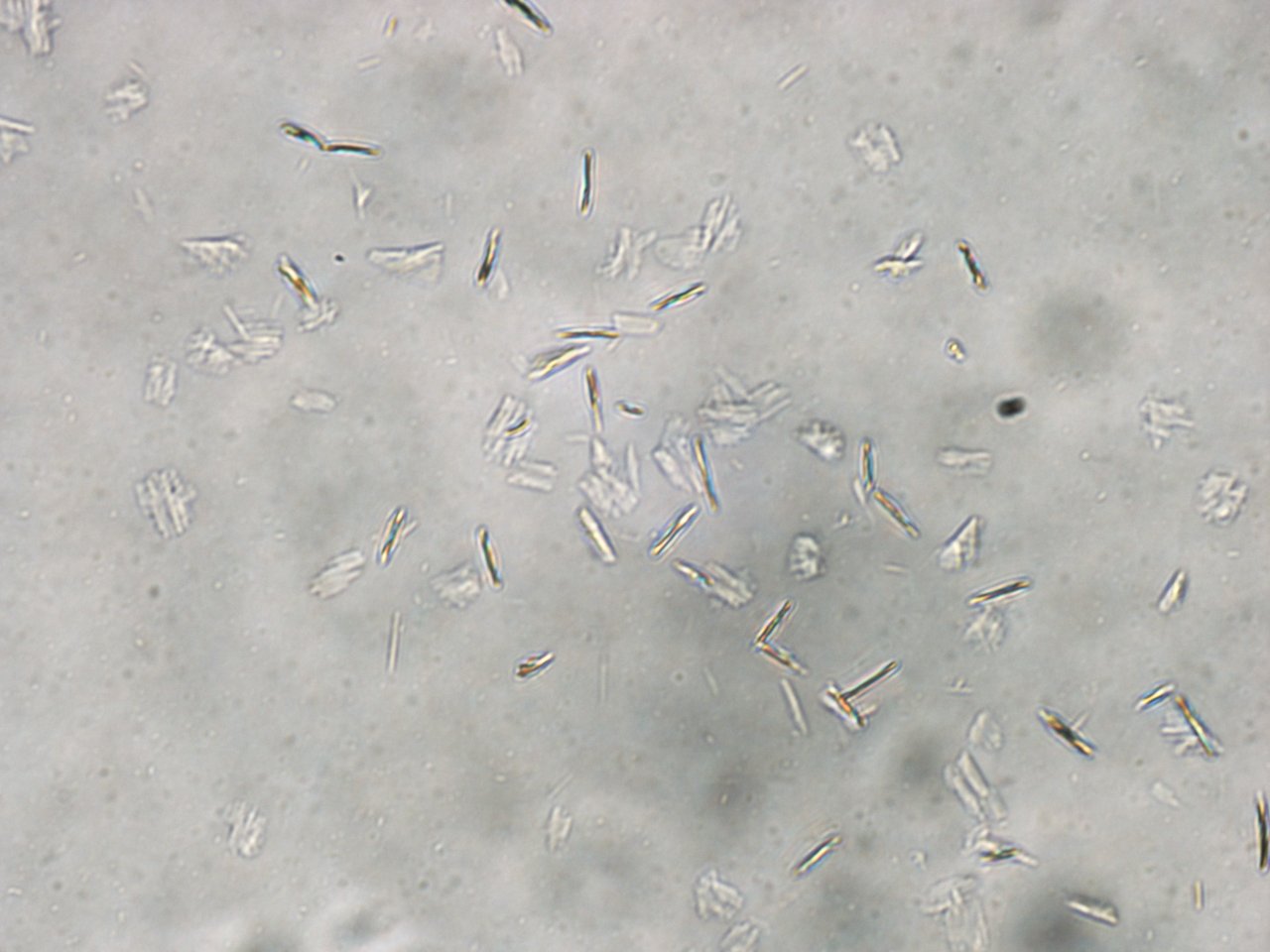

Supporting Figure 4: Polarized light microscope image of GI R387A crystals (10min, 100x objective).

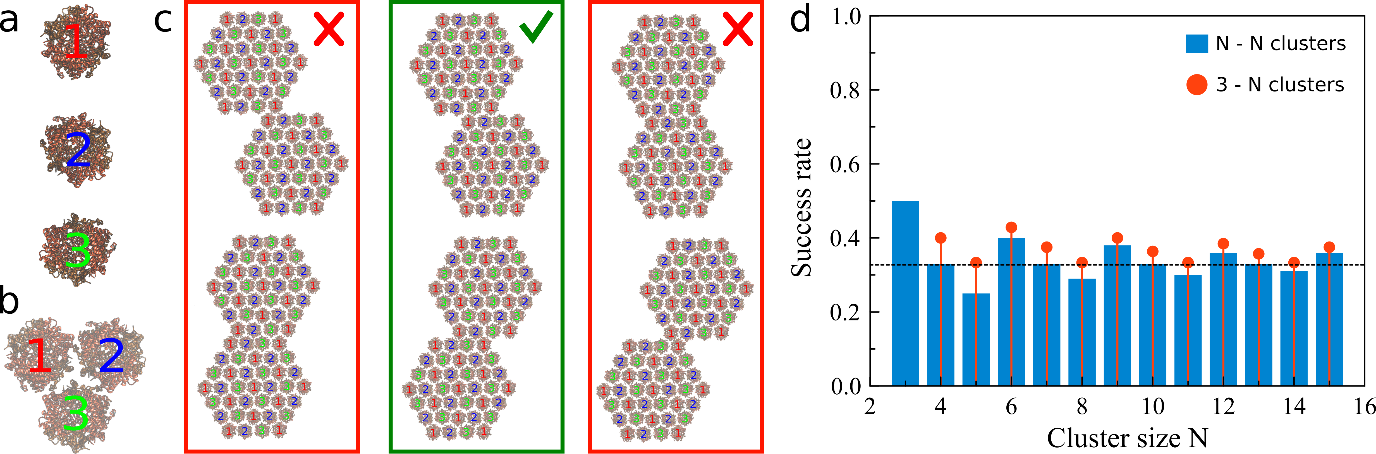

Supporting Figure 5: (a) The three different molecular orientations of a GI tetramer within the (001) plane of the H32 space group; (b) local arrangement around the three fold screw axis; (c) possible configurations for two clusters with facets of size N=4 to interact laterally: 2 out of 6 configurations lead to succesfull bond formation; (d) probabilities for succesful docking derived from the configurations in (b) as a function of cluster size, where N indicates the number of molecules for each facet: N-N denotes interactions between two clusters of size N, whereas 3-N corresponds to docking of a cluster of size 3, with clusters of varying size N.

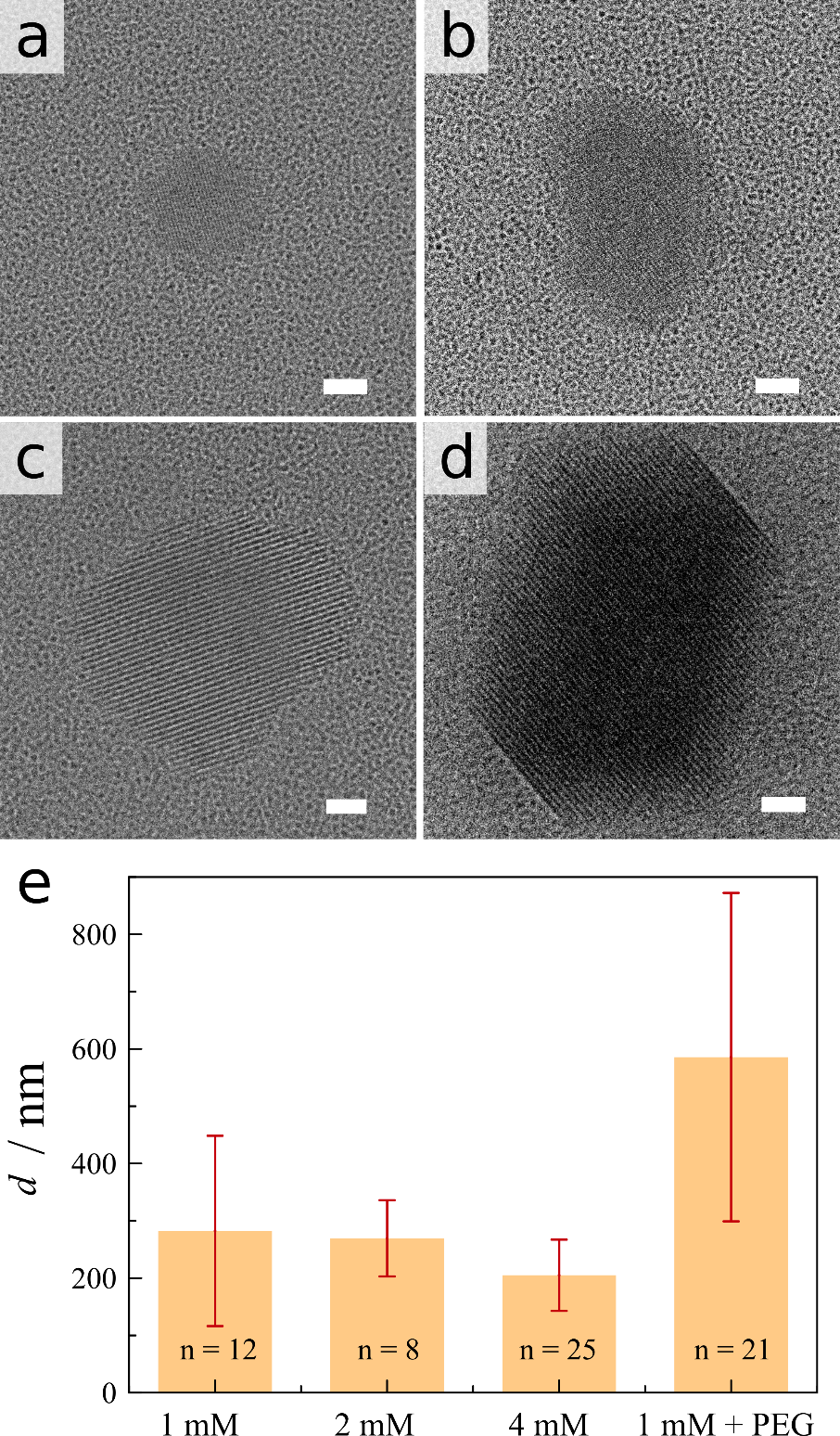

Supporting Figure 6: (a)-(d) I222 GI nanocrystals detected in concentrated glucose isomerase solutions at 173 mg.mL^-1^, 10mM Hepes pH 7.0, 1mM MgCl_2_. Scalebar is 50nm; (e) Typical nanocrystal dimension is independent (within error) on MgCl_2_ for 1mM, 2mM and 4mM MgCl_2_ but increases after addition of PEG-1000.

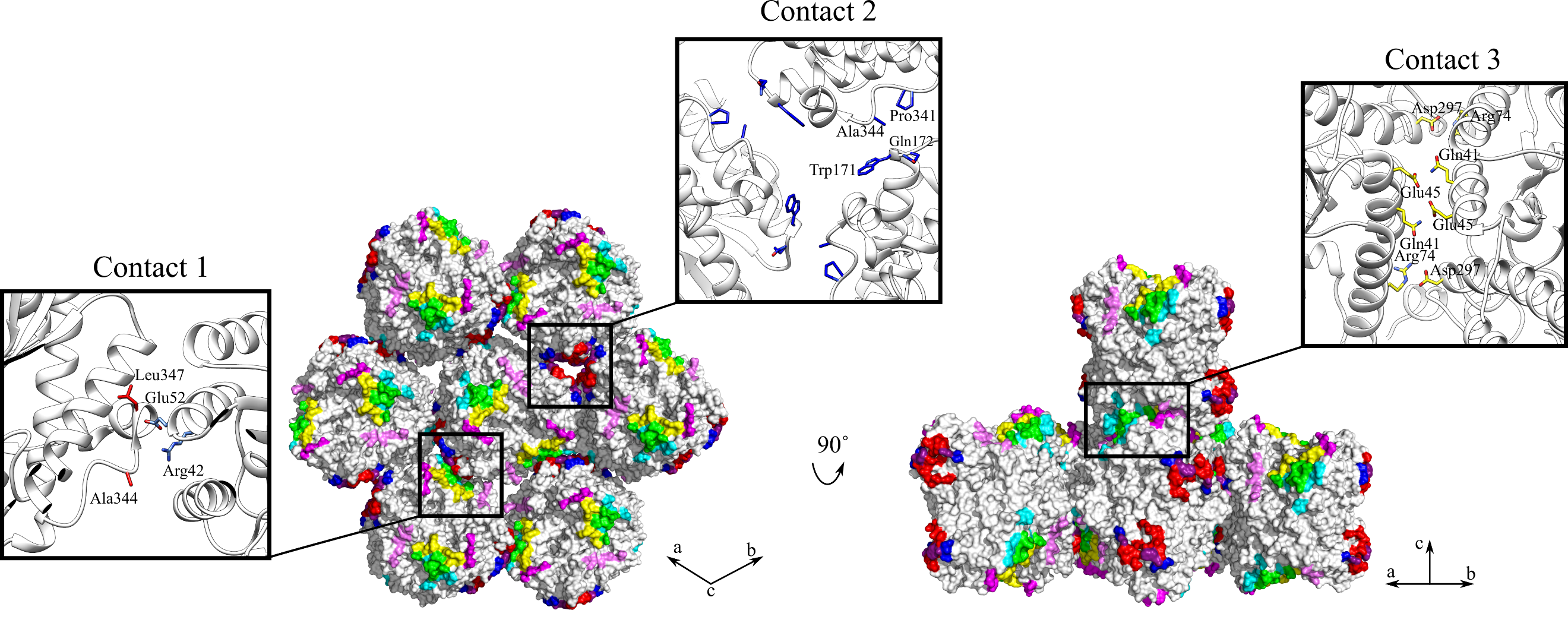

Supporting Fig.7: Surface rendering of the nearest neighbors and crystal lattice contacts of glucose isomerase in the trigonal space group H32: residues that partake in crystal packing contacts are colored according to the color scheme shown in Supporting Table 1. Note that some residues are involved in two different contacts; the corresponding regional overlaps between neighboring patches are color coded: i.e. patch H1a (red), H2 (blue), H1a/H2 overlap (purple); H1b (cyan), H3 (yellow), H1b/H3 overlap (green).

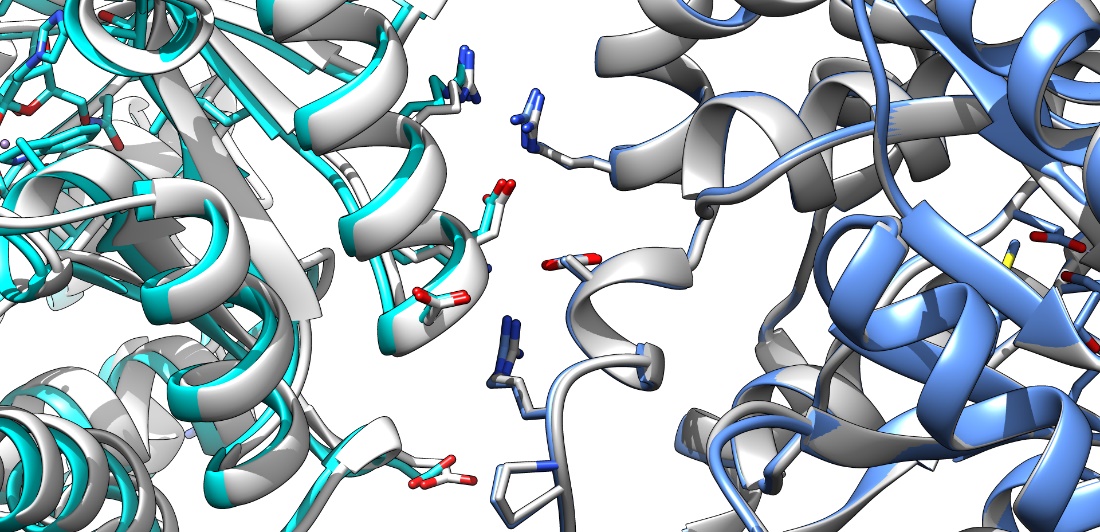

Supporting Figure 8: Comparison of the lattice contact for the I222 space group grown without Mg^2+^ (symmetry related molecules shown in white and grey) and with Mg^2+^ (cyan and blue). The former contact is based on the GI structure with PDB id 9XIA, which was crystallized in 10mM PIPES 7.4, 0.76M (NH4)_2_SO_4_ and 25 mg.mL^-1^ glucose isomerase, whereas the latter is based on a GI structure that was determined in-house by diffracting crystals that were grown in 10mM Hepes 7.0, 20mM MgCl_2_ and 30 mg.mL^-1^ glucose isomerase.

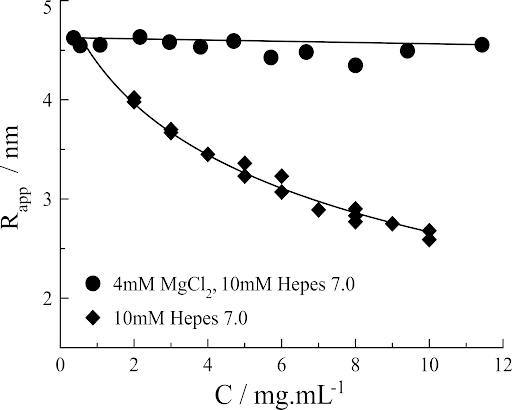

Supporting Figure 9: Apparent hydrodynamic radius of GI determined using DLS, as a function of protein concentration at 0mM and 4mM MgCl_2_. Lines are guides for the eye.

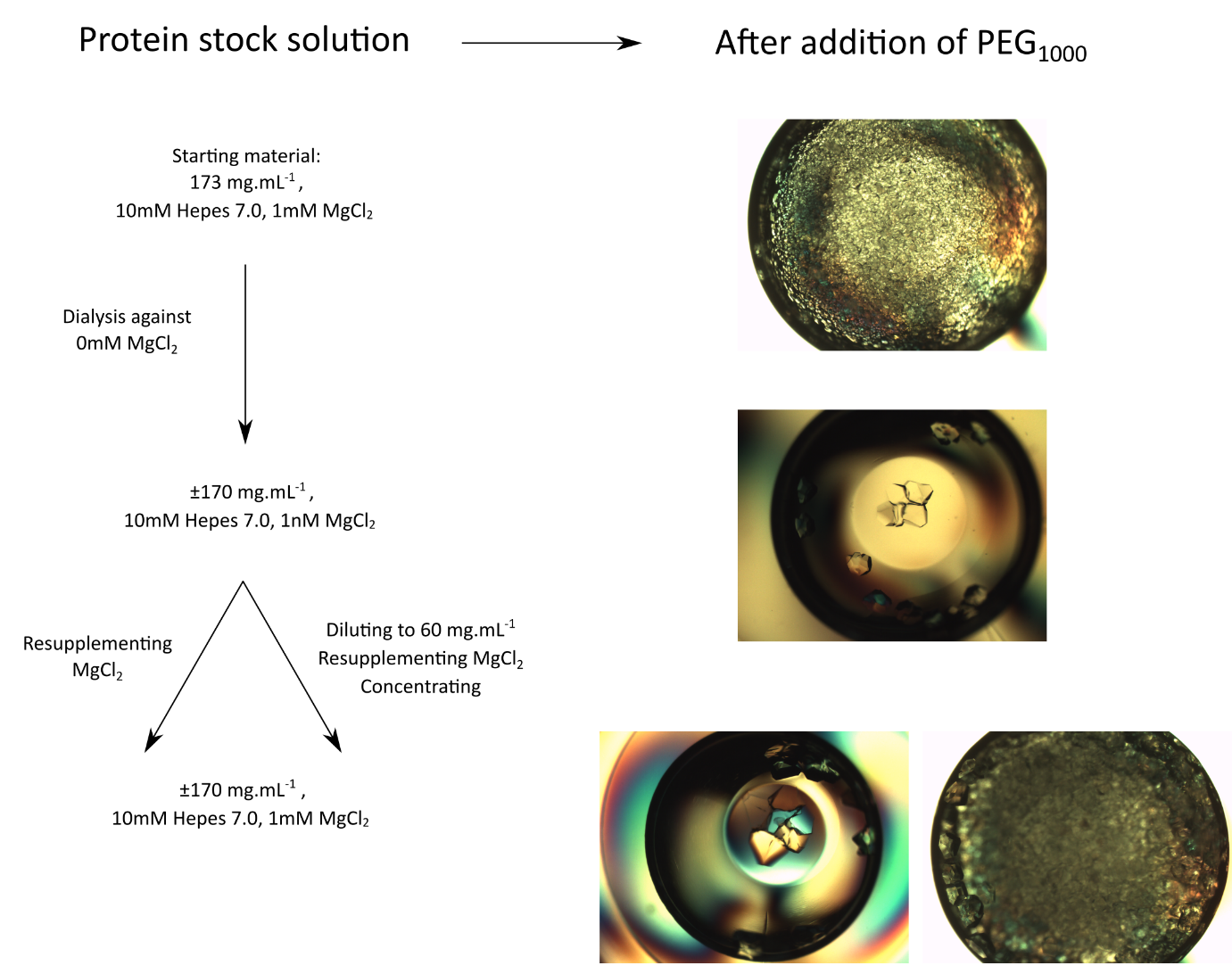

Supporting Figure 10: Reversibility of the seeding effect upon removal and subsequent addition of MgCl_2_ during the concentrating step.

Supporting Discussion

**Lattice contact analysis of H32 R387A crystals**

Here we evaluate the molecular lattice contacts within the H32 space group. As we do not have crystallographic data for macroscopic R387A H32 crystals (due to twinning), we base our analysis on diffraction data that was collected for H32 crystals that were grown from a closely related point mutant, dubbed S171W GI. This approach is valid because there is good agreement between the unit cells derived from X-ray data for S171W (Supporting Table 2) and those derived from the cryoEM data for R387A (see Supporting Fig.2). In the H32 space group, each GI molecule has eight nearest neighbors (Supporting Fig.7). Six of these lie within the (001) plane and involve the formation of two different types of lattice contacts, C1 and C2 (Supporting Table 1). The remaining two neighbors are formed along the c-axis each making the same C3 contact with the central molecule. These crystallographic contacts (C1, C2 and C3) are formed by the recognition of different patches on the molecular surface of GI. C1 involves the burial of patch H1a on the central reference molecule and H1b on the corresponding symmetry mate. C2, however, is formed by the self-recognition of a single surface patch, denoted as H2. This means that two identical regions on the surfaces of two symmetry related copies interface to produce contact C2. Similarly, the contact C3 comprises two identical copies of patch H3, but also the interfacing of two minor patches H3a and H3b. It is also important to note that each GI molecule is a homo-tetramer with C2 symmetry, which means that there are multiple copies of each patch on a single tetramer. The H32 contacts described above are unique for this space group, there is little to no overlap in terms of the associated surface patches (I1a and I1b^7^) with the contacts that exist in I222 crystals. This may explain why we do not see any GI H32 nanocrystals that serve as an epitaxial platform for the formation of an I222 lattice, and indicates that nucleation of each polymorph occurs independent of the other.

**The presumed pre-nucleation GI particles are I222 nanocrystals.**

Based on our cryo-EM results we conclude that the mesoscopic particles that exist in concentrated GI stock solutions are I222 nanocrystals that are structurally equivalent to the macroscopic I222 crystals we detect in typical crystallization experiments. This readily explains the seeding behavior at the macroscopic level, as well as the specificity regarding the I222 polymorph. But it also raises certain questions regarding their origin, stability and nanoscopic size.

We addressed their stability by trying to ascertain if the GI stock solution is in equilibrium or supersaturated with respect to the I222 phase; undersaturation is not expected as it would be a thermodynamic violation. For this we determine the crystal-liquid coexistence curve as a function of the concentration of MgCl_2_ (main text, Fig.5b). MgCl_2_ is typically added to GI in low millimolar concentrations because the active site holds two metal cofactors^32^ (e.g. Co, Mn, Mg) that contribute to the thermal stability of the protein^33^ but it can also affect the solubility^34^. As it turns out, the GI equilibrium concentration for the conditions used here is, highly sensitive to [MgCl_2_], decreasing 500-fold from 1mM to 20mM MgCl_2_. Note that there is no salting-in regime, a common feature in the low salt region of the phase diagram of many proteins; rather GI exhibits only salting-out over the tested range. Such high sensitivity to MgCl_2_ could indicate that Mg^2+^ interacts in a specific manner with the surface of GI, or indeed in the formation of salt bridges in lattice contacts, as has been seen for other protein crystals (e.g. insulin, Zn^2+^, PDB: 1G7A; ferritin, Cd^2+^, PDB: 1LB3). To this end, we collected an in-house X-ray dataset on I222 crystals grown in 20mM MgCl_2_ but found no meaningful differences with the lattice contact of I222 crystals grown in 0.8M ammonium sulfate (Supporting Fig.8; Supporting Table 2). Moreover, crystallization is not specific to Mg^2+^, we obtain qualitatively similar results with other divalent cations as well, such as Ca^2+^, Zn^2+^, Ni^2+^, Cu^2+^, Mn^2+^ which all trigger I222 nucleation in the 2.5 to 50mM range. In fact, monovalent cations can also be used as a salting-out agent, albeit at slightly higher concentrations (e.g. 100mM NaCl). It therefore seems likely that the cations we tested facilitate macromolecular self-assembly by shielding the electrostatic repulsion between GI molecules in a non-specific manner (at pH 7.0, GI has a net negative charge of -74e^35^). This is reflected in the apparent hydrodynamic radius (*R_app_*) of the monomers measured by DLS (Supporting Fig.9). In 10mM Hepes 7.0, *R_app_* decreases monotonically as a function of the GI concentration. The slope (*k_D_^-1^*) of this curve is a measure of the pairwise interaction potential between the GI molecules averaged over all possible orientations^36^. The negative sign of *k_D_^-1^* indicates net repulsion between the solute molecules. *k_D_^-1^* approaches zero for the curve collected at 4mM MgCl_2_, implying that the repulsive forces between the particles are greatly diminished, which is in line with the general trend of the solubility curve.

Based on our solubility measurements we conclude that the crystals in main text Figure 5 are not in equilibrium with their surrounding liquid: the bulk GI concentration was 173 mg.mL^-1^, which is 6% higher than the equilibrium concentration recorded for this condition (C_e_ is 163 mg.mL^-1^ at 1mM MgCl_2_). Since the mother liquor is still supersaturated, we expect that the nanocrystals can continue to grow until equilibrium is reached. Such an equilibrium may not necessarily be reached in a practical timeline given that the kinetics could be very slow as a result of an additional barrier that traps these nanocrystals in a metastable state. Clues to the origins of such a barrier can be found in the crystal morphology (main text Fig.5a, Supporting Figure 6a-d). The crystal facets are often straight with sharp corners at the edges, suggesting full completion of the outer layers. From this we conclude that crystals with fully developed habits are of (near) magic size. Magic sized clusters are metastable towards further expansion because new molecular layers need to be initiated on the existing facets for growth to continue. It is the work associated with the formation of a new molecular island that gives rise to the metastability. We can estimate the impact on the kinetics of crystal growth by predicting the two-dimensional nucleation rate using kinetic parameters that we determined previously^37,38^. For a supersaturation of ln(163/173)=0.059, we predict a nucleation rate of two-dimensional islands to be 3.6 x 10^9^ m^-2^s^-1^. Assuming nucleation to be the rate-limiting step, this translates roughly into the deposition of one new molecular layer per day. Such a slow growth rate fits indeed with the typical crystal sizes that we observe in cryoTEM.

If we lower the MgCl_2_ concentration by dialyzing the 1mM condition to 10mM Hepes 7.0 (expected final [MgCl_2_] = 1nM), then the seeding effect upon mixing with 8% (w/v) PEG 1000 is lost, i.e. we see a drastic reduction of the final crystal count towards the level we observe for a filtered solution (Supporting Fig.10). This suggests that the nanocrystals have dissolved because of the undersaturation that was imposed. If we resupplement that solution with MgCl_2_ to a final concentration of 1mM than we do not see an increase in the final crystal count, which means that nanocrystals likely not reformed. This brings us to the final conundrum: why did nanocrystals form in our initial stock solutions (173 mg.mL^-1^ GI, 1mM MgCl_2_) in the first place? The answer can be found in the last preparation step of the protein stock solution where we concentrate the dialyzed GI (typically 60 mg.mL^-1^) via a spin-concentrator. We found that there is a noticeable build-up of GI at the bottom of the concentrator. We recorded GI concentrations in *ex situ* aliquots that are in the order of 300 mg.mL^-1^ but higher local concentrations (and therefore supersaturations) likely exist *in situ* as well. Note that these gradients are removed at the end of the concentrating step by mixing the entire volume. Now, if we resupplement MgCl_2_ prior to concentrating to the solution from which MgCl_2_ was removed via dialysis, then we indeed recover a drastic increase in the number of crystals detected after concentrating and mixing with PEG, demonstrating full reversibility.

| Contact | H | S | Patch | Symmetry | ΔASA | Residues |
| --- | --- | --- | --- | --- | --- | --- |
| C1 | 2 | 0 | H1a | X,Y,Z | 390 | K159, L164, E167, T170, W171, E207, R208, **A344**, D345, G346, **L347**, Q348, A349 |
|  |  |  | H1b | -Y-1,X-Y-1,Z | 384 | Q4, P5, E38, Q41, **R42**, A44, **E45**, D81, T82, D297, W300 |
| C2 | 0 | 0 | H2 | -X,-X+Y,-Z | 59 | W171, Q172, G173, P341, A344 |
| C3 | 2 | 3 | H3 | X-Y-1/3,-Y-2/3,-Z+1/3 | 350 | R32, D35, V37, E38, **Q41**, R42, **E45**, R74, D295, F296, D297 |
|  |  |  | H3a | X,Y,Z | 88 | E67, E70, K73 |
|  |  |  | H3b | -X+Y-1/3,-X-2/3,Z+1/3 | 83 | E328, A332, R334, A364, R368 |

Supporting Table 1: Lattice-contact analysis of glucose isomerase in space group H32: two contacts (C1 and C2) are made within the (001) plane, and one contact (C3) is made along the *c*-axis. The columns ‘H’ and ‘S’ denote the number of hydrogen bonds and salt-bridges involved in the respective contacts. The ΔASA is the change in surface accessible area in Å^2^ of the surface patches that are involved in the contact formation. Residues that are involved in a hydrogen bond or salt bridge with a nearest-neighbor residue are shown in bold and red, respectively. Residues highlighted in purple or green are shared between different patches (purple: H1a and H2; green: H1b and H3).

Supporting Table 2: X-ray crystallography: data collection and refinement statistics.

|  | **I222 (20mM MgCl_2_)** | **H32 (S171W)** |
| --- | --- | --- |
| Resolution (Å)^a^ | 2.04 (2.04–2.1) | 2.13 (2.13–2.19) |
| Space group | *I222* | *H32* |
| Cell dimensions (Å; a, b, c); angle (°; α, β, γ) | 92.6, 97.9, 102.3;  90, 90, 90 | 132.9, 132.9, 234.9;  90, 90, 120 |
| Total/Unique reflections | 203,460/ 29,534 | 605,407/ 44,769 |
| Completeness (%)^a^ | 99.2 (95.0) | 99.9 (99.5) |
| *R*_merge_^a^ | 0.038 (0.089) | 0.145 (1.49) |
| *R*_pim_^a^ | 0.023 (0.060) | 0.059 (0.620) |
| I/σ(I)^a^ | 43.0 (22.6) | 11.2 (1.7) |
| CC_1/2_ | 0.999 (0.995) | 0.998 (0.562) |
| Multiplicity | 6.9 (5.3) | 13.5 (12.9) |
| *R_cryst_* | 15.40% | 19.60% |
| *R*_free_ | 18.50% | 22.80% |
| Rmsd in bond lengths (Å) | 0.011 | 0.012 |
| Rmsd in bond angles (°) | 1.746 | 1.80 |
| B- factor statistics (Å^2^) | | |
| Protein all atoms | 29.40 | 36.81 |
| Protein main chain atoms | 27.43 | 34.81 |
| Protein side chain atoms | 31.40 | 38.86 |
| Mn ions | 33.52 | 78.96 |
| Solvent atoms | 32.55 | 38.93 |
| Ramachandran statistics (Molprobity) | | |
| Favoured | 96.20% | 95.45% |
| Outliers | 0.20% | 0.26% |

^a^Values in parentheses refer to the highest resolution shell.
